# A novel, NADH-dependent acrylate reductase in *Vibrio harveyi*

**DOI:** 10.1101/2022.01.20.477176

**Authors:** Yulia V. Bertsova, Marina V. Serebryakova, Alexander A. Baykov, Alexander V. Bogachev

## Abstract

Bacteria coping with oxygen deficiency use alternative terminal electron acceptors for NADH regeneration, particularly fumarate. Fumarate is reduced by the *FAD_binding_2* domain of cytoplasmic fumarate reductase in many bacteria. The variability of the primary structure of this domain in homologous proteins suggests the existence of reducing activities with different specificities. Here we produced and characterized one such protein, *Vibrio harveyi* ARD, and found it to be a specific NADH:acrylate oxidoreductase. This previously unknown enzyme contains covalently bound FMN and non-covalently bound FAD and FMN in a ratio of 1:1:1. The covalently bound FMN is absolutely required for activity and is attached by the specific flavin transferase, ApbE, to a threonine residue in the auxiliary *FMN_bind* domain. RT-qPCR and activity measurements indicated dramatic stimulation of ARD biosynthesis by acrylate in the *V. harveyi* cells grown aerobically. In contrast, the *ard* gene expression in the cells grown anaerobically was high without acrylate and increased only twofold in its presence. These findings suggest that the principal role of ARD in *Vibrio* is energy-saving detoxification of acrylate coming from the environment.

**Importance:** The benefits of the massive genomic information accumulated in recent years for biological sciences have been limited by the lack of data on the function of most gene products. Approximately half of the known prokaryotic genes are annotated as “proteins with unknown functions,” and many other genes are annotated incorrectly. Thus, the functional and structural characterization of the products of such genes, including identification of all existing enzymatic activities, is a pressing issue in modern biochemistry. In this work, we have shown that the *ard* gene product of *V. harveyi* (GenBank ID: AIV07243) exhibits a yet undescribed NADH:acrylate oxidoreductase activity. This activity may allow acrylate detoxification and its use as a terminal electron acceptor in anaerobic or substrate in aerobic respiration of marine and other bacteria.

## INTRODUCTION

Many microbes are capable of anaerobic respiration by using various organic and inorganic compounds instead of oxygen as terminal electron acceptors in the respiratory electron transport chain. Fumarate is a typical organic terminal acceptor commonly reduced by membrane-bound quinol:fumarate oxidoreductase complex, similar to the Complex II of the respiratory chain (1). In some microbes, fumarate is reduced by a soluble monomeric enzyme using cytochrome *c*, NADH, or flavin as the source of reducing equivalents during anaerobic respiration (2–6). In both types of fumarate-reducing enzymes, fumarate is bound and converted by a *FAD_binding_2* domain (Pfam ID: PF00890).

Soluble cytoplasmic NADH:fumarate oxidoreductase from the facultative anaerobic bacterium *Klebsiella pneumoniae* (FRD) is formed by three domains: *OYE-like, FMN_bind*, and *FAD_binding_2* (5, 7) (Fig. 1A). The *FAD_binding_2* domain contains a fumarate-binding site and a non-covalently bound FAD molecule. The *OYE-like* domain (PF00724) contains a non-covalently bound FMN molecule and is responsible for NADH oxidation. The domain *FMN_bind* (PF04205) carries out electron transfer between the two other domains and contains an FMN residue covalently linked through a phosphoester bond (5). The catalytic mechanism of fumarate reduction has been described for the soluble fumarate reductase (flavocytochrome *c*, Fcc_3_) of the bacterial genus *Shewanella* (2, 3). The reaction involves hydride ion transfer from a nearby (3.2 Å) FAD molecule to the fumarate C2 atom, accompanied by proton transfer to the C3 atom via a proton-conducting channel, and yields succinate (3). X-ray structural studies of *Shewanella frigidimarina* Fcc_3_ have pinpointed three groups of functionally important amino acid residues: those binding the C4 carboxylate (Arg402, His504, and Arg544; Motif I) and C1 carboxylate (His365 and Thr377; Motif II) and those forming the proton conductance channel (Glu378, Arg381, and Arg402; Motif III) (Fig. 1B) (3). All these residues are conserved in *K. pneumoniae* fumarate reductase (Table 1).

**Fig. 1.**
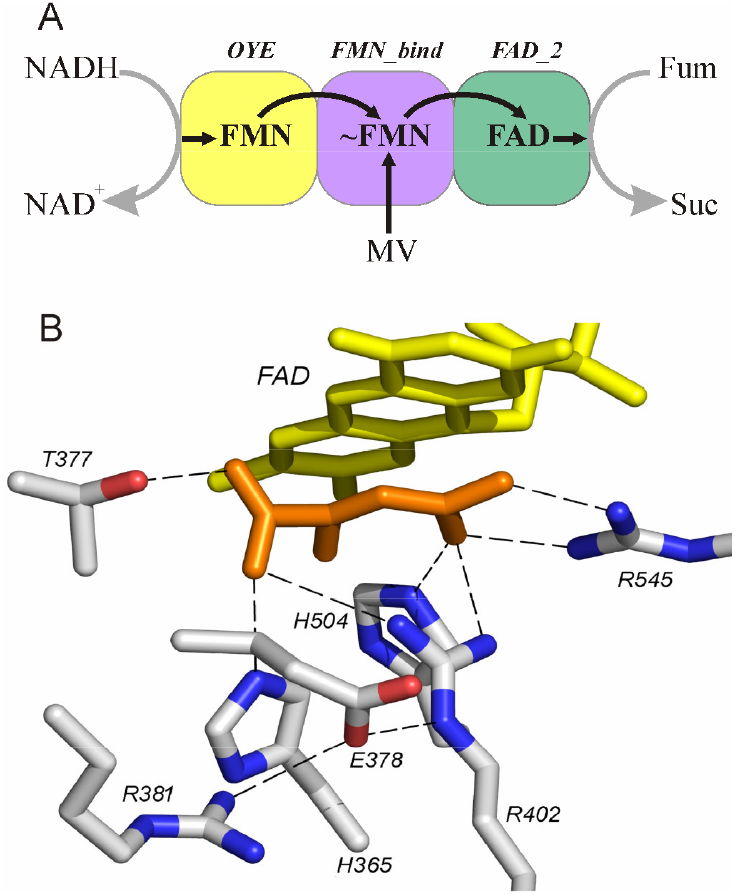
The structural aspects of fumarate reductase. (A) The domain artichecture, prosthetic groups, and electron transport pathways from NADH/MV to fumarate in *K. pneumoniae* FRD. ~FMN denotes covalently bound FMN. (B) The structure of the substrate-binding site in the *FAD_binding_2* domain of *S. frigidimarina* Fcc_3_ with bound malate molecule, shown in orange (PDB ID: 1QJD). Dashed lines show the polar contacts that fix the positions of malate carboxylates.

**Table 1.** Enzymatic activities and conserved amino acid residues of *FAD_binding_2* domain-containing proteins

| Protein source | GeneBank<br>protein ID | Pfam domains | Substrate<br>reduced | Partial alignment of the *FAD_binding_2* domains |  |  |
| --- | --- | --- | --- | --- | --- | --- |
|  |  |  |  | Motif 1 | Motif II | Motif III |
| *S. frigidimarina* | ABI73517 | *FAD_binding_2*<br>*Cytochrom_c3_2* | Fumarate | Arg402...His504...Arg544 | His365...Thr377 | Glu378...Arg381...Arg402 |
| *K. pneumoniae* | BAH63234 | *FAD_binding_2*<br>*FMN_bind*<br>*OYE-like* | Fumarate | Arg756...His859...Arg899 | His719...Ser731 | Glu732...Arg735...Arg756 |
| *Geobacter*<br>*sulfurreducens* | AJY71931 | *FAD_binding_2*<br>*Cytochrom_c3_2* * | Methacrylate | Arg353...His461...Arg501 | Tyr311...Val323 | Glu324...Arg330...Arg353 |
| *Wolinella*<br>*succinogenes* | CAE09288 | *FAD_binding_2*<br>*Cytochrom_c3_2* * | Methacrylate | Arg344...His454...Arg495 | Gly301...Val313 | Ala387...Arg390...Arg411 |
| *S. oneidensis*<br>MR-1 | Q8CVD0 | *FAD_binding_2*<br>*FMN_bind* | Urocanate | Arg411...His520...Arg560 | Tyr373...Ile386.. | Ala387...Arg390...Arg411 |
| *V. harveyi* | AIV07243 | *FAD_binding_2*<br>*FMN_bind*<br>*OYE-like* | ? | Arg834...His940...Arg980 | Met796...Thr808 | Gly809...Val812...Arg834 |

\*This domain is present as an additional subunit. All other listed proteins are monomeric.

Genes encoding proteins with the *FAD_binding_2* domain in their structures are ubiquitous among bacterial genomes. However, some of these proteins are unlikely to be fumarate reductases because the residues responsible for C1 carboxylate binding and proton transfer are not conserved in this domain. Three of such proteins have been characterized and indeed found to exhibit different substrate specificities—they preferentially reduce methacrylate (8, 9) or urocanate (10, 11) with cytochrome *c* as the second substrate (Table 1). Motifs I and III are highly conserved in all these enzymes, whereas Motif II is only partially conserved. These findings suggest that substrate specificity within this group of fumarate analogs is determined by Motif II residues, consistent with the X-ray crystallographic data for *S. frigidimarina* fumarate reductase (3).

Here we tested this prediction with the *FAD_binding_2* domain-containing protein ARD (GenBank ID: AIV07243) found in the *Vibrio harveyi* genome. ARD and similar proteins of the genus *Vibrio* resemble *K. pneumoniae* NADH:fumarate oxidoreductase by domain composition but largely differ from this enzyme and all other reductases listed in Table 1 not only in Motif II but also in Motif III (Table 1). Specifically, the histidine residue of Motif II is replaced by methionine, and only one arginine residue (affiliated with both Motifs I and III) is conserved in Motif III in comparison with the fumarate reductase sequences. Consistent with these structural differences, the results below identified ARD as the reductase enzyme with yet unknown substrate specificity.

## RESULTS

### 1. Isolation of *V. harveyi* ARD and identification of bound flavins

The *ard* gene was amplified from *V. harveyi* genomic DNA and cloned into the pBAD-TOPO vector that adds a 6×His tag to protein C-terminus. The ARD *FMN_bind* domain contains a D_449_VISGA**T**_455_ signature motif for the attachment of FMN via a covalent phosphoester bond to the threonine residue (12). To verify this prediction, we expressed the *ard* gene in *E. coli* in the absence and presence of the plasmid pΔhis3 (13), which harbors the gene for the cytoplasmic form of *V. cholerae* flavin transferase ApbE with broad substrate specificity (12). Metal affinity chromatography of the cytoplasmic fractions of the ApbE-lacking and ApbE-producing cells yielded bright yellow 95-kDa proteins (Fig. 2B) in both cases (the calculated mass of 6×His tagged ARD is 112 kDa). A MALDI-MS analysis of the trypsin-digested protein bands revealed the full-length *ard* gene product in the ARD preparations isolated from ApbE-containing and ApbE-lacking cells (sequence coverage of 86 and 92%, respectively). Furthermore, both the N- and C-terminal peptides were present in the mass spectrum, and their amino acid sequences were confirmed by MS/MS analysis (see Supplementary Material). The difference between the measured and theoretical masses is not thus caused by ARD fragmentation and likely results from its anomalous (increased) electrophoretic mobility.

**Fig. 2.**
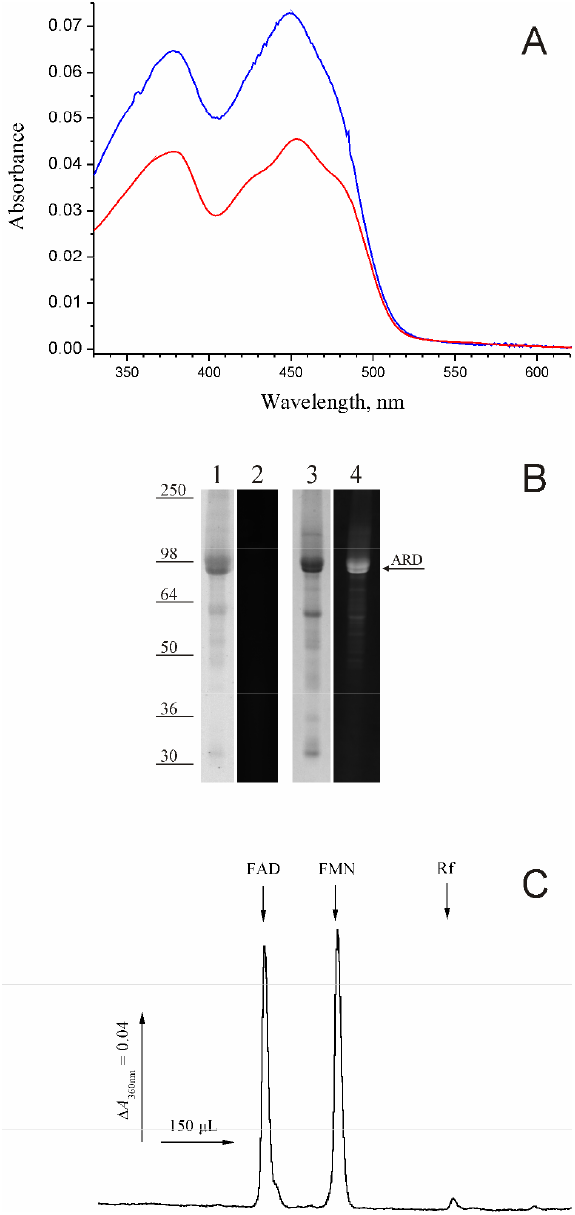
Identification of the bound flavin groups. (A) Electronic absorption spectra of ARD preparations (1.8 μM) purified from ApbE-lacking (red curve) or ApbE-containing (blue curve) *E. coli* strain. Spectra were determined in 10 mM Tris-HCl buffer, pH 8.0, containing 100 mM KCl. (B) SDS-PAGE of ARD isolated from ApbE-lacking (lanes *1* and *2*) or ApbE-containing (lanes *3* and *4*) *E. coli* cells. The gels were stained with Coomassie Blue (lanes *1* and *3*) or photographed under illumination at 473 nm without staining (lanes *2* and *4*). The protein load was 4 μg per lane. The bars with numbers on the left side denote positions and molecular masses of marker proteins. (C) HPLC separation of non-covalently bound flavins of ARD. Retention times for authentic FAD, FMN, and riboflavin (Rf) are indicated by arrows.

The optical spectra of the isolated proteins (Fig. 2A) demonstrated absorbance maxima at ~380 and ~450 nm, characteristic of flavin prosthetic groups. The absorbances were lower with ARD isolated from ApbE-lacking cells. Moreover, this protein did not fluoresce after SDS-PAGE when illuminated at 473 nm, unlike the protein isolated from ApbE-producing cells (Fig. 2B). These observations indicated covalently and non-covalently bound flavin groups in authentic ARD, the latter group(s) being attached by the flavin transferase ApbE.

To identify the non-covalently bound flavins, the flavinylated ARD was denatured and separated from the solution by ultrafiltration through a 50-kDa cut-off filter. The filtrate contained two-thirds of the initial flavin content, as determined spectrophotometrically. HPLC analysis of the filtrate identified FAD and FMN in equal amounts (Fig. 2C).

To identify the flavinylation site, we subjected the ARD bands in the PAGE gel (lanes *1* and *3* in Fig. 2B) to in-gel trypsin digestion and MALDI-MS and MS/MS analysis. Based on ApbE specificity (12), the flavinylation site was predicted in the ARD *FMN_bind* domain within the tryptic peptide I436PQQILDGQTLN<u>IDVISGA**T**</u>VSSQAVLDGVSNAVDLAGGNSEALR480 (the flavinylation motif is underlined, the acceptor residue shown in bold). This peptide should demonstrate the monoisotopic MH^+^ masses of 4944.5 in the flavinylated and 4506.3 in the non-flavinylated states. The mass spectrum of the tryptic digestion of ARD produced in the ApbE-lacking *E. coli* cells demonstrated only the presence of the non-flavinylated peptide (Fig. 3B), whereas the flavinylated peptide predominated in the spectrum of ARD produced by the *ard*-*apb*E coexpression (Fig. 3A). The MS/MS analysis of the peptide with the *m/z* of 4944.5 (Fig. 3B) revealed two main signals with *m/z* of 4488.3 and 4506.3 in the spectrum of fragmentation. A likely corollary is that they resulted from FMN loss in the full (*m/z* = 456) or dehydrated (*m/z* = 438) form (14). Other than *m/z* = 4944.5 candidates for flavinylated peptides were not observed in the tryptic digest of ARD.

**Fig. 3.**
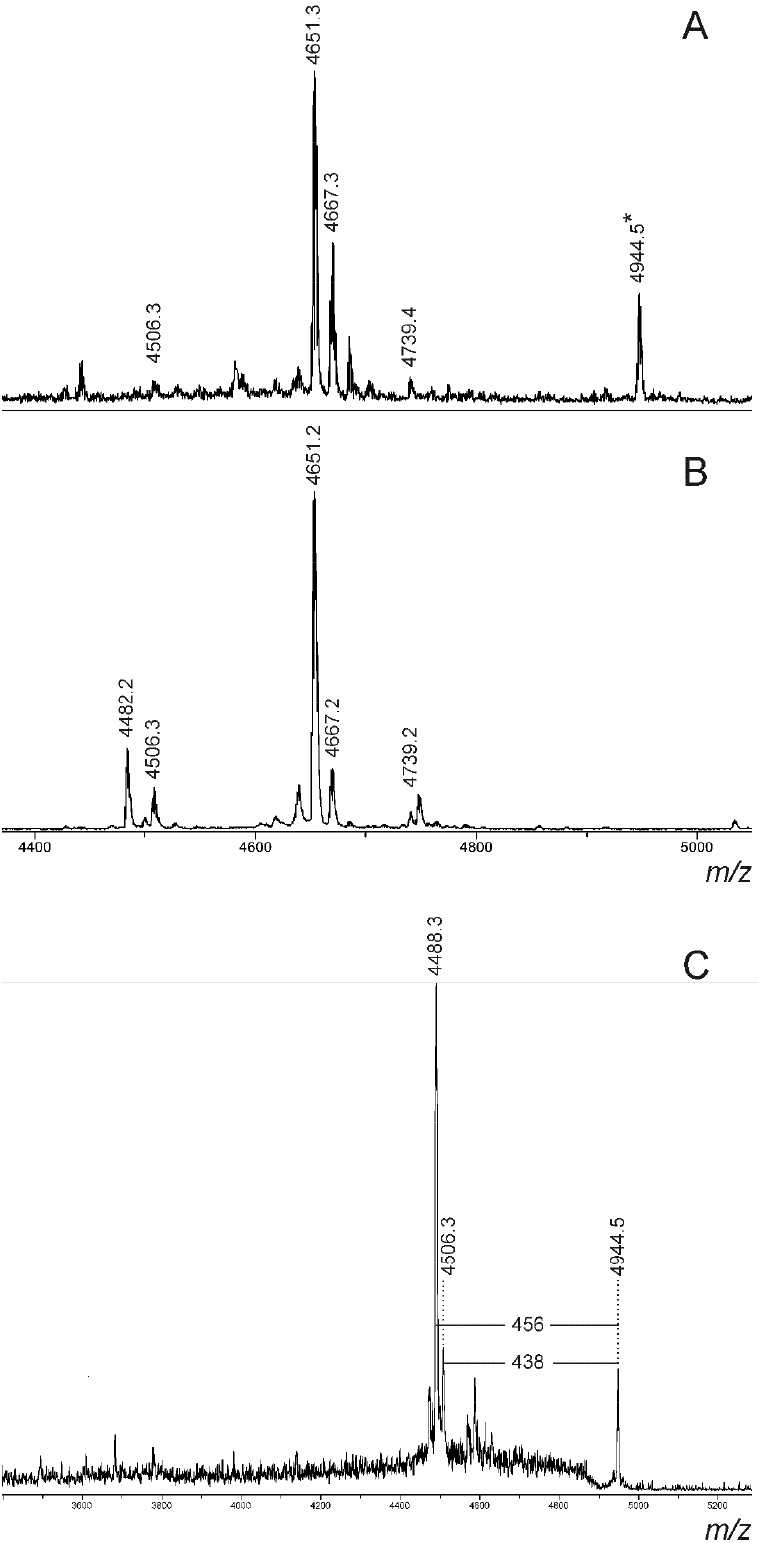
Parts of the MALDI mass spectra of in-gel tryptic digests of ARD produced in ApbE-containing (A) or ApbE-lacking (B) *E. coli* strain. The asterisk on panel A marks the peak of the flavinylated peptide *(m/z* of 4944.5). (C) MS/MS spectrum of the peptide with *m/z* of 4944.5.

Our data indicate thus that ARD resembles *K. pneumoniae* cytoplasmic NADH:fumarate oxidoreductase in that both proteins contain covalently bound FMN and non-covalently bound FAD and FMN in a ratio 1:1:1. Based on these data, the spectra of ARD in the native state and after denaturation with SDS (1%, 5-min incubation at 80°C), and the known extinction coefficients for FAD and FMN in solution (15), the extinction coefficient for native ARD at 450 nm was estimated to be 39.5 mM^-1^ cm^-1^. This estimation was based on the premise that denaturation brings flavins into an aqueous environment.

### 2. The enzymatic activities of ARD

The substrate specificity of ARD was tested with a series of α,β-unsaturated carbonic acids, listed in Table 2, as tentative electron acceptors from methyl viologen (MV). A negligible or minimal activity was observed with most tested compounds, except for methacrylate and, in particular, acrylate. The carboxyl group in the substrate is necessary, as no MV oxidation was detected when acrylamide was used as an electron acceptor. The reactions with acrylate and methacrylate were explored in some detail below.

**Table 2.**
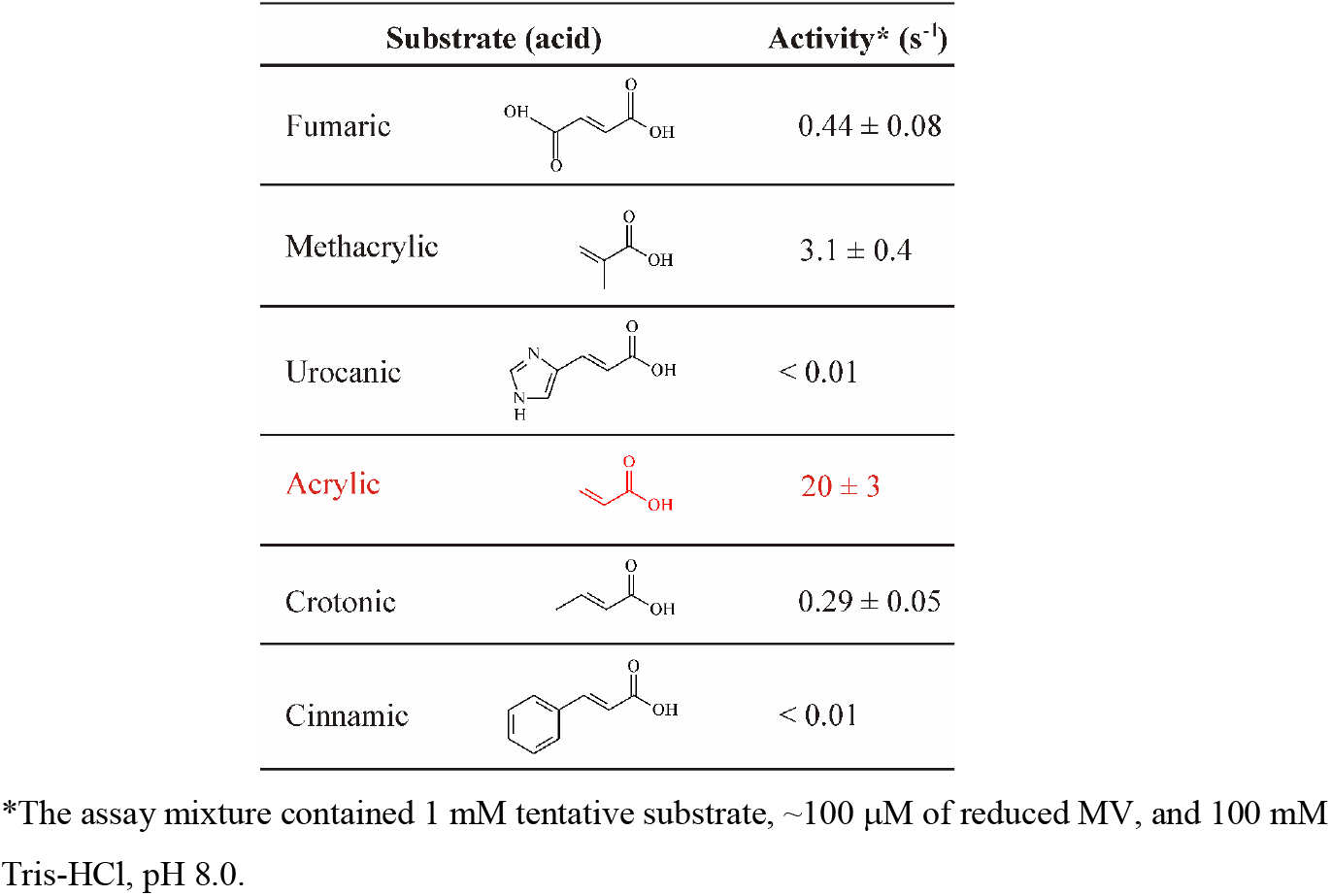
The reductase activity of ARD against a series of α,β-unsaturated carbonic acids measured with MV as the electron donor.

| Substrate (acid) | | Activity* ($s^{-1}$) |
| --- | --- | --- |
| Fumaric | | $0.44 \pm 0.08$ |
| Methacrylic | | $3.1 \pm 0.4$ |
| Urocanic | | $< 0.01$ |
| Acrylic | | $20 \pm 3$ |
| Crotonic | | $0.29 \pm 0.05$ |
| Cinnamic | | $< 0.01$ |

\*The assay mixture contained 1 mM tentative substrate, $\sim 100 \mu M$ of reduced MV, and 100 mM Tris-HCl, pH 8.0.

The similarity of the domain structures of ARD and *K. pneumoniae* NADH:fumarate oxidoreductase, in particular the presence of an *OYE-like* domain in both, raised the possibility of NADH being the physiological electron donor for ARD (5). Indeed, NADH supported acrylate and methacrylate reduction, as illustrated in Fig. 4 for acrylate. Acrylate was absolutely required for NADH oxidation under anaerobic conditions and markedly accelerated the reaction under aerobic conditions, wherein acrylate competed with oxygen as the alternative electron acceptor. Importantly, acrylate did not support NADH oxidation by ARD deficient in covalently bound flavin (Fig. 4, *upper trace*).

**Fig. 4.**
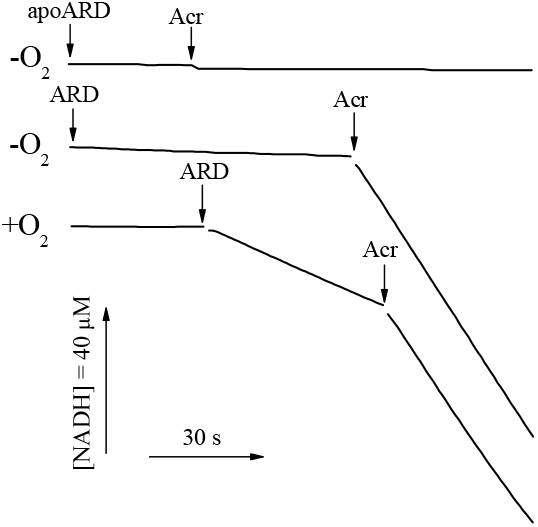
Typical traces of NADH oxidation by ARD under anaerobic (-O_2_) or aerobic (+O_2_) conditions. ARD isolated from the ApbE-lacking *E. coli* strain is designated as apoARD. Arrows indicate additions of 30 nM ARD or apoARD and 1 mM acrylate (Acr). The assay mixture contained 150 μM NADH and 100 mM Mes-KOH, pH 6.5. The anaerobic samples were also supplemented with 10 mM glucose, 5 U/mL glucose oxidase, and 5 U/mL catalase.

The NADH:acrylate oxidoreductase activity was pH-dependent, with a maximum between pH 6.5 and 7.0 (see Supplementary Material). The apparent *K*_m_ value for NADH measured at pH 6.5 with 1 mM acrylate was 14 ± 1 μM. NADPH could not replace NADH as the electron donor. Steady-state kinetic analysis was also performed at varied acrylate and methacrylate concentrations using a fixed saturating NADH concentration (150 μM). This analysis yielded the *K*_m_ values of 16 ± 1 for acrylate and 690 ± 140 μM for methacrylate. The respective *k*_cat_ values for these substrates were 19 ± 1 and 2.5 ± 0.2 s^-1^. Acrylate is thus a much “better” substrate for ARD.

HPLC identified the sole product of acrylate reduction by ARD to be propionate (Fig. 5). Thus, the ARD-catalyzed oxidoreductase reaction can be described as follows:

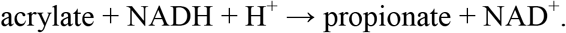

**Fig. 5.**
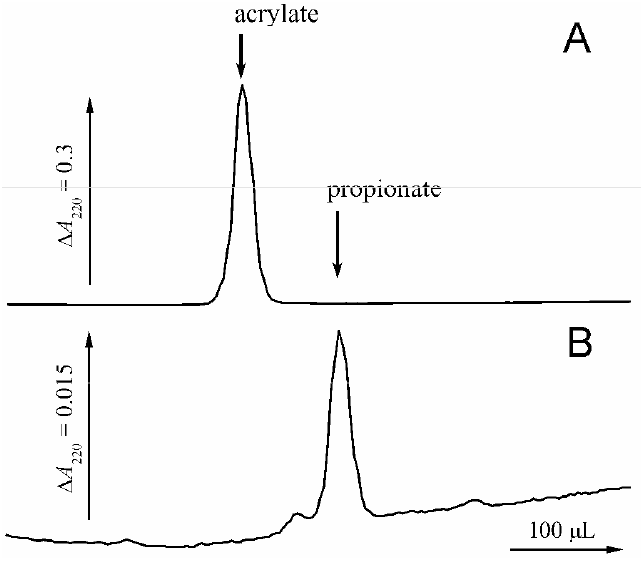
HPLC identification of the product of acrylate reduction by ARD. Acrylate (5 mM) was reacted with 20 mM sodium dithionite and 50 μM MV in the presence of 0.5 μM ARD. (A) The extract of the reaction mixture before ARD addition. (B) The same after a 60-min incubation with ARD. The elution volumes for authentic acrylic and propionic acids are indicated by arrows, the corresponding retention times were 184 and 211 s. Of note, that extinction coefficients for acrylate and propionate at 220 nm differed 20-fold.

ARD did not catalyze the reverse reaction of propionate oxidation with phenazine methosulfate or dichlorophenolindophenol as the electron acceptor, resembling in this aspect all known soluble fumarate reductases and related enzymes (2).

### 3. Dependence of ARD biosynthesis in *V. harveyi* cells on growth conditions

An RT-qPCR study indicated that the level of *ard* expression was quite low in the *V. harveyi* cells grown aerobically in the absence of acrylate but increased ~100-fold in the presence of 2 mM acrylate, the maximal concentration at which acrylate does not yet inhibit cell growth significantly (Table 3). In contrast, the expression of *ard* was high without acrylate and increased only twofold in its presence when the cells were grown under anaerobic conditions. The maximal level of *ard* expression was higher under anaerobic conditions. The enzymatic activity of ARD in cell lysates demonstrated similar trends (Table 3). These results indicated that either oxygen absence or acrylate presence induces ARD biosynthesis. The effects of the two factors are additive, and *ard* is maximally expressed under anaerobic conditions in the presence of acrylate.

**Table 3.** Correlation between *ard* transcription level and the acrylate reductase activity in the *V. harveyi* cells cultivated in different conditions.

| Growth conditions | *ard* mRNA/rRNA $\times 10^{-6}$* | ARD activity (nmol·min$^{-1}$ ·mg$^{-1}$)* |
| --- | --- | --- |
| +O<sub>2</sub>, no acrylate | 0.094 ± 0.022 | Not detected (<2) |
| +O<sub>2</sub>, 2 mM acrylate | 9.3 ± 0.8 | 18 ± 4 |
| -O<sub>2</sub>, no acrylate | 7.5 ± 0.2 | 10 ± 4 |
| -O<sub>2</sub>, 2 mM acrylate | 14.2 ± 1.1 | 38 ± 7 |

\*From three independent experiments.

## DISCUSSION

*V. harveyi* ARD contains a Met residue instead of His in the position that determines fumarate C1 carboxylate binding in fumarate reductases (Table 1) and was therefore expected to have a different substrate specificity. The results reported above indicated that ARD and possibly other similar enzymes possess a previously unknown acrylate reductase activity. This is the principal activity of ARD, which converts alternative substrates at much lower rates (Table 2). The bulky methionine residue in the binding site of ARD appears to be the critical factor that determines ARD specificity. Only fumarate and its analogs with smaller substituents in the left part are converted at measurable rates, whereas those with more bulky substituents are not converted. Methacrylate is a better alternative substrate than crotonate (Table 2), indicating that the α-position in acrylate is more tolerable to substitutions than the β-position. The significant activity against the bulkier fumarate may be explained by the conservation in ARD of the threonine residue (Thr808) found, together with histidine, in Motif II of fumarate reductases (Table 1).

Although ARD can reduce methacrylate, its catalytic efficiency (*k*_cat_/*K*_m_) with this substrate is smaller by a factor of 330 compared to acrylate. Oppositely, known methacrylate reductases do not reduce acrylate (16) or reduce it at much lower rates than methacrylate (8, 9). The two enzymes also differ markedly in the active site motifs (Table 1).

Acrylate was a potent inducer of gene *ard* transcription, under aerobic conditions in particular (Table 3), providing independent support to the notion that acrylate reductase is the principal activity of ARD. On the other hand, these data shed light on the possible physiological role of ARD in *V. harveyi* cells. The ability of acrylate to markedly induce ARD production under both aerobic and anaerobic conditions suggests a role for ARD in acrylate detoxification. Acrylate toxicity results from its being a potent electrophile capable of reacting with many cellular components (17, 18). A known way of acrylate detoxification involves NADPH-dependent reduction of acryloyl-CoA by the protein AcuI to yield propionyl-CoA (18). However, acryloyl-CoA is much more toxic than acrylate (18), making ARD-catalyzed acrylate removal before its conversion to acryloyl-CoA a more efficient way of detoxification. In these circumstances, the role of AcuI may be auxiliary and consist in neutralizing acryloyl-CoA formed from acrylate in competing reactions before its conversion by ARD. The proposed role of ARD in acrylate detoxification is consistent with the gene *ard* localization in close vicinity to the gene for another detoxicating enzyme, glutathione S-transferase (AIV07248 in *V. harveyi*), on chromosome 2 in most *ard*-containing *Vibrio* species.

The high level of *ard* expression under anaerobic conditions even in the absence of acrylate (Table 3) suggests that ARD may use acrylate as the terminal electron acceptor for NADH regeneration at oxygen deficiency. The ability of *Halodesulfovibrio aestuarii* (formerly known as *Desulfovibrio acrylicus*) to use acrylate as terminal electron acceptor has also been documented, although the enzyme(s) responsible for the acrylate reduction has(ve) not been identified (19). Moreover, *V. harveyi* cells can use acrylate as the only carbon and energy source (20). This makes acrylate detoxification in *V. harveyi* an energy-saving process—instead of consuming energy, like in the detoxification pathways involving glutathione S-transferase, the ARD-catalyzed reaction produces propionate, a “fuel” under aerobic conditions. Also of importance, acrylate reduction helps deplete cells of NADH, which is crucial under anaerobic conditions. The interplay between ARD’s detoxification and energy production functions is determined by oxygen availability. As *ard* expression is induced by acrylate under both aerobic and anaerobic conditions, detoxification seems to be the primary function. If NADH reoxidation were the primary function of ARD, the induction would have been only observed in the absence of oxygen, like with other enzymes, such as NADH:fumarate oxidoreductase (5) and urocanate reductase (10). Further, if the primary function were propionate accumulation, ARD production would be stimulated by acrylate only under aerobic conditions because propionate is not fermentable.

The exogenous nature of acrylate in *V. harveyi* is supported by the findings that this bacterium is a common inhabitant of acrylate-rich niches, for instance, coral mucus (20). Marine bacteria of the *Vibrio* genus demonstrate a positive taxis test with acrylate (21). What is the source of acrylate in the environment? The primary precursor of acrylate in the living world is dimethylsulfoniopropionate (DMSP) (22), which is produced and accumulated in high concentrations in many marine algae as a compatible solute. DMSP conversion into acrylate is catalyzed by the DMSP lyases DddL, DddP, DddQ, DddW, and DddY (22), explaining the abundance of acrylate in many marine ecological niches and the presence of ARD homologs in the *Vibrio* bacteria, which often inhabit these niches. Noteworthy, genes for the DMSP lyases are absent in the *V. harveyi* genome, and *V. harveyi* cells do not convert DMSP or use it as a sole carbon source (data not shown), which is consistent with the exogenous nature of acrylate in them.

Acrylate reductase activity appears to be also indispensable in many terrestrial niches, including human symbionts and pathogens. Genes for ARD-like proteins are found in the genomes of certain non-marine bacteria, such as *Staphylococcus pasteuri* and *S. warneri* (human pathogens); *S. lugdunensis*, *S. massiliensis*, and *S. epidermidis* (humane skin microbiome); *Dickeya zeae* and *Paenibacillus brasilensis* (plant pathogens and symbionts); *Gilliamella apicola* (bee intestine symbiont). The occurrence of ARD-containing bacteria in human pathogens may make ARD an important drug target. Disturbances in the gut microbiome may cause the accumulation of the toxic products of fermentation and anaerobic respiration, leading to various diseases (23). The case of urocanate respiration is relevant and illustrative in this respect. Molinaro *et al*. (24) demonstrated an association of urocanate respiration with type 2 diabetes, and Koh *et al.* (25) further determined that imidazolylpropionate, the product of urocanate reduction, causes glucose intolerance by inhibiting mTORC1 and, hence, suppressing insulin signal transduction. These findings, which opened new perspectives in type 2 diabetes treatment, emphasize the importance of identifying alternative anaerobic respiration types in microbes.

## MATERIALS AND METHODS

### Production and isolation of ARD

An expression vector for the C-terminal 6×His-tagged ARD protein was constructed by amplifying the *ard* gene (GenBank Gene ID: 57822953) by PCR with a high fidelity Tersus PCR kit (Evrogen) and the primer pair 5’-GAGGAATAAATTTATGGCTCAGTTAGTCGAT / 5’-GGAAACAAGATCAGATGCACTCTC using the genomic DNA of *V. harveyi* as the template. The resulting 3028-bp fragment was cloned into the pBAD-TOPO vector (Invitrogen), yielding the pBAD_VHRD plasmid. The 6×His-tagged ARD protein was produced in *Escherichia coli* BW25113 or /pΔhis3 cells and purified using metal chelate chromatography as described previously (7).

### Analysis of flavin groups

The relative amounts of covalently and non-covalently bound flavins in the recombinant ARD were determined by HPLC and spectral analyses of the denatured protein as described by Barquera *et al.* (26). Non-covalently bound flavins were separated by HPLC as described elsewhere (7). SDS-PAGE was performed using 10% (w/v) polyacrylamide gels (27). The gels were stained with PageBlue™ protein staining solution (Fermentas). Covalently bound flavins were detected by scanning unstained gels with a Typhoon™ FLA 9500 laser scanner (GE Healthcare) with excitation at 473 nm and the detection of emission using the SYBR Green II protocol according to the manufacturer’s recommendations. The in-gel trypsin digestion of ARD and MALDI-MS and MS/MS analysis were done as described previously (14).

### Enzymatic activities

NADH- and MV-supported activities of ARD were determined spectrophotometrically by following the oxidation of NADH or MV (5). The assay medium contained 1 mM MV, 1 mM electron acceptor, and 100 mM Tris-HCl (pH 8.0) or 150 μM NADH, 1 mM electron acceptor, 10 mM glucose, 5 U/mL glucose oxidase, 5 U/mL catalase, and 100 mM Mes-KOH (pH 6.5). MV was pre-reduced with sodium dithionite until the absorbance at 606 nm of approximately 1.5 was obtained. The NADH-supported reaction was also measured under aerobic conditions without glucose oxidase and catalase. One unit of enzyme activity was defined as the amount of the enzyme that catalyzes the oxidation of 1 μmol NADH or 2 μmol MV per 1 min.

The pH-dependence of NADH:acrylate oxidoreductase activity was measured in a 100 mM BIS-TRIS propane-citrate buffer, pH 4.5–9.0, containing 5 U/mL glucose oxidase, 5 U/mL catalase, 10 mM glucose, 150 μM NADH, and 1 mM potassium acrylate.

In the experiments aimed at identification of the product of acrylate reduction, the reaction medium contained 5 mM acrylate, 20 mM sodium dithionite, 50 μM MV, and 100 mM Tris-HCl (pH 8.0). The reaction was performed in a 3.2-mL sealed cuvette completely filled with the reaction mixture. Acrylate reduction was initiated by adding 0.5 μM ARD and terminated after 60 min by adding 5% (vol./vol.) HClO_4_. In the control experiment, HClO_4_ was added before ARD. Precipitated protein was removed by centrifugation, and carbonic acids were isolated from the samples by double extraction with diethyl ether (2 × 1.5 mL). The ether was evaporated under a stream of air, and the residue was dissolved in 1 mL of aqueous 0.05% H_3_PO_4_. The samples thus prepared were separated by HPLC on a ProntoSil-120-5-C18 AQ column using a Milichrom A-02 chromatograph (both from Econova, Novosibirsk, Russia). The column was pre-equilibrated with aqueous 0.05% H_3_PO_4_ solution and eluted with a linear gradient of 0% to 10% aqueous methanol containing 0.05% H_3_PO_4_ at a flow rate of 0.2 mL/min with UV detection at 220 nm.

Propionate dehydrogenase activity was assayed by measuring the reduction of 2,6-dichlorophenolindophenol (DCPIP) at 600 nm. The assay mixture contained 2 mM Tris-propionate, 2 mM phenazine methosulphate, 25 μM DCPIP, and 100 mM Mes-KOH, pH 6.5. *Cultivation of V. harveyi cells*

*V. harveyi* cells were grown aerobically or anaerobically at 32°C up to the middle of exponential phase in a mineral medium containing 20 g/L NaCl, 0.7 g/L KCl, 1 g/L MgSO_4_×7(H_2_O), 0.5 g/L NH_4_Cl, 0.5 mM Na_2_HPO_4_, 2 g/L sucrose, 0.5 g/L yeast extract, 50 mM Tris-HCl (pH 8.0). When required, the medium was supplemented with 2 mM potassium acrylate. Cleared cell lysate was prepared as described (28) and assayed for MV:acrylate oxidoreductase activity. Protein concentration was determined by the bicinchoninic acid method (29) using bovine serum albumin as the standard.

### Quantitative reverse transcription polymerase chain reaction (RT-qPCR)

RNA extraction from *V. harveyi* cells and cDNA synthesis were performed as described (30). RT-qPCR assays were done with the qPCRmix-HS SYBR kit (Evrogen), using the cDNA preparations as templates and 5’-CGGCGCAAACACATACCT / 5’-GCCCGGTTGTTCAATCTCT as primer pair. 16S rRNA was used for data normalization (primer pair 5’-CAGCCACACTGGAACTGAGA / 5’-GTTAGCCGGTGCTTCTTC). Serial dilutions of *V. harveyi* genomic DNA, wherein the genes for ARD and 16S rRNA are present in a 1:12 ratio (31), were used for calibration.

## Acknowledgements

This work was supported by the Russian Science Foundation (project # 22-24-00133). MALDI MS and laser scanner facilities became available to us in the framework of the Moscow State University Development Program PNG 5.13.

