## Supplementary material for "A novel, NADH-dependent acrylate reductase in *Vibrio harveyi*": Supplementary_Material.pdf

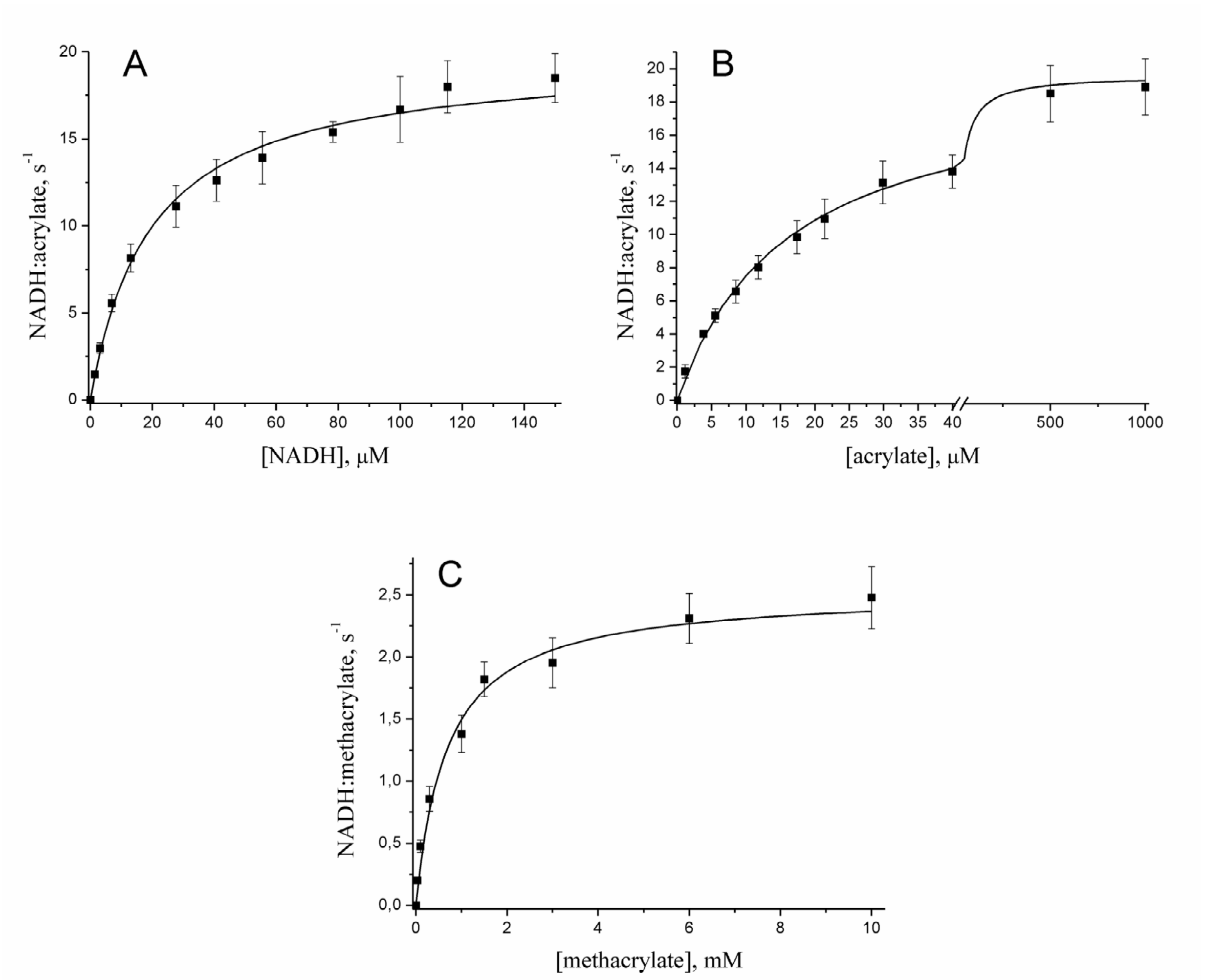


**Fig. S1.** Dependence of the NADH:acrylate oxidoreductase activity of ARD on the concentrations of NADH (A), acrylate (B), and methacrylate (C). The lines show the best fits of the Michaelis-Menten equation.

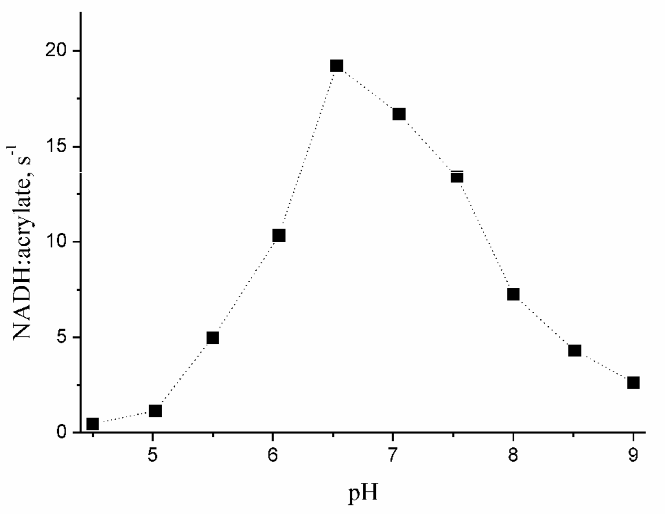


**Fig. S2.** Dependence of the NADH:acrylate oxidoreductase activity of ARD on pH.

The supplementary material also includes separate files with the results of the MALDI-MS and MS/MS analyses of ARD isolated from ApbE-containing (Mascot_ARD_msms.mht) and ApbE-lacking cells (Mascot_apoARD_msms.mht).
